# Antidiabetic action of Mcy protein through the regulation of number and affinity of insulin receptors

**DOI:** 10.1101/2021.01.22.427879

**Authors:** Saritha Marella, Peddanna Kotha, S. Abdul Nabi, B.P. Girish, Kameswara Rao Badri, Apparao Chippada

## Abstract

Evidence based immunological cross-reactivity studies and anti-diabetic investigations have suggested the presence of insulin-like peptides in plants. Mcy protein, isolated from the fruits of *Momordicacymbalaria*, was shown to have antihyperglycemic, antihyperlipidemic activities along with renal as well as hepatoprotective activities in Streptozotocin induced diabetic rats.Mcy protein was shown to have insulin like and/or insulin secretagoguestructure and/or functions. Hence, the present study was conducted to elucidate molecular mechanism wherebyMcy protein elicits its therapeutic role and also to know whether the Mcyprotein has any structural and functional similarity with insulin. Results of our experiments revealed that the Mcyprotein is insulin like protein. Further, we assessed the effect of treatment of Mcy protein on the levels of glucose transport (glucose transporter (GLUT-2) and on the levels of key regulators of glucose and lipid metabolisms like hepatic glucokinase (GK) and sterol regulated element binding protein-1c (SREBP-1c). Our findings demonstrated that Mcy protein decreased the elevated expressions of GK, SREBP-1c and GLUT-2 that were observed in diabetic animals. Insulin-receptor binding studies using rat erythrocytes demonstrated that mean specific binding of insulin with insulin receptors was significantly increased in Mcy treated diabetic rats when compared to diabetic control rats. Scatchard analyses of insulin-binding studies yielded curvilinear plots, and the number of receptor sites per cell was found to be 180±21.1 in Mcy treated diabetic animals calculated to be significantly superior to that of diabetic control animals. Kinetic analyses also revealed an increase in the average receptor affinity of erythrocytes from Mcy treated rats compared to diabetic control rats suggesting acute alteration in the number and affinity of insulin receptors on the membranes of erythrocytes.

## Introduction

Various insulin like proteins were reported from plants. The seminal investigation on the presence of insulin-like antigens was published by Khanna et al. (1976) who isolated and patented insulin like active principle called p-insulin from the fruits of *Momordica charantia*. Recently, Mcy protein was isolated by our group from the fruits of *Momordica cymbalaria* and reported to possess significant antidiabetic activities including antihyperglycemic and antihyperlipidemic activities (Saritha et al., 2016; Marella et al., 2015). Partial sequence of Mcy protein and insulin level determination studies in Mcy protein treated animals have shown insulin like and/or insulin secretagogue activity of Mcy protein (Saritha et al., 2016; Marella et al., 2015). However, structural details and molecular mechanism of action of Mcy protein is elusive.

To understand the mechanism of action of Mcy protein, we studied the levels of key regulatory factors that regulate glucose metabolism and lipid metabolism like glucose transport and the levels of glucokinase (GK) and sterol regulatory element binding protein (SREBP)-1c. Glucose homeostasis is regulated in liver, an insulin sensitive tissue, through dynamic interactions between glucose utilization and production pathways. Thus, a potent anti-diabetic agent should improve glucose induced insulin secretion, hepatic glucose metabolism, and peripheral insulin sensitivity (Ferre et al., 1996). In mammals, glucokinase, the major glucose phosphorylating enzyme expressed in hepatocytes and pancreatic β-cells, plays a critical role in maintaining the glucose homeostasis (Matschinsky et al., 1993). Long-term regulation of hepatic GK activity is controlled through its expression in liver and is reliant on the presence of insulin (Bourbonias et al., 2012). Other critical group of crucial factors that regulate glucose metabolism, SREBPs, belong to helix-loop-helix leucine zipper family of transcription factors that regulate fatty acid and cholesterol synthesis (Brown and Goldstein, 1997; 1999). SREBP-1c is a key player in carbohydrate and lipid metabolisms. Regardless of all the intracellular regulation/s, glucose transport is the first and rate limiting step (Maughana, 2009) in carbohydrate metabolism, which is facilitated by glucose transporters (GLUT-2) across the cell membrane (Anand et al., 2010). Drugs which facilitate GLUT-2 translocation and improve insulin sensitivity together with carbohydrate and lipid metabolism could be beneficial for the treatment of diabetes (Kipmen-Korgun et al., 2009).

The first and major check point of insulin action *in vivo* is its binding to receptors wherein this biological process is subjected to modulation by alterations in either the receptor number and/or affinity (De Pirro et al., 1980). The rat erythrocytes serve as a valuable, accurate, sensitive, and efficient model system for preclinical receptor studies. Many studies related to insulin binding to its receptors have been demonstrated in tissues of different animal species wherein proven to decrease the binding in diabetes mellitus (Olefsky and Kolterman, 1981). However, with the introduction of human erythrocyte insulin receptor assay by Gambhir et al. (1977) erythrocytes were studied to possess insulin binding characteristics which help in studying the role of phytoconstituents in the treatment of diabetes mellitus (Murugan et al., 2008). The status of insulin receptors and the receptor binding of erythrocytes from diabetic subjects/models were used to evaluate therapeutic potential of new pharmaceuticals by evaluating their ability to compete with radio labelled ligands (Pari et al., 2004). Hence, we elucidated the binding of insulin to its receptors in Mcy treated and untreated rat erythrocytes to shed light on the mechanism of action of Mcy protein.

## Materials & Methods

### Isolation and purification of Mcy protein

Mcy protein was isolated from the fruits of *Momordica cymbalaria* as described earlier (Saritha et al., 2016; Marella et al., 2015).

### Mcy protein: insulin like molecule

The electrophoretic separation of protein was done by discontinuous SDS-PAGE according to Laemmli et al. (1970). The proteins separated on SDS–PAGE were transferred to nitrocellulose membrane using western blot unit (MerckMillipore, India). Western blotting was carried out following standard protocol using anti-insulin (mouse 1:1000dilution, Sigma, USA), donkey anti-mouse (1:5000 dilution). Blot was developed by ECL prime reagents (GE healthcare) using versa doc (BioRad). The PVDF membrane was stained with Ponceau’s stain followed by destaining with TBST.

### Induction of diabetes mellitus and Mcy protein treatment

Male Wistar rats weighing 180-200 g were used for these studies as per institutional (Sri Venkateswara University, Tirupati, India) and international guidelines (Saritha et al., 2016; Marella et al., 2015).

To verify the potential hypoglycemic activity and to confirm the anti-hyperglycemic activity of Mcy protein experiments were designed as outlined below.

Two groups of normal rats (6 rats in each group) were treated with either Mcy protein or insulin as described below. Insulin treated group was used as control and Mcy protein dose was used as determined by us earlier (Saritha et al., 2016; Marella et al., 2015).

Group I: Normal rats treated with 1.9mg/kg bw of Mcy protein [NMT].

Group II: Normal rats treated with 2.5U/kg bw of Insulin (Huminsulin 40IU/ml) [NIT].

Two groups of experimental induced diabetic rats were treated as described below. Diabetes was induced using 50mg/kg body weight of streptozotocin as described earlier [2,3]. Blood glucose levels greater than or equal to 250mg/dl were considered diabetic and used for the current study.

Group I: Diabetic rats treated with 1.9mg of Mcy protein/kg bw [DMT].

Group II: Diabetic rats treated with 2.5U of Insulin/kg bw [DIT].

The blood glucose levels were monitored for every 30min by glucose oxidase peroxidase method (Saritha et al., 2016; Marella et al., 2015). To prevent hypoglycemic shock in the insulin treated animals, glucose solution was given orally as soon as the hypoglycemia was observed.

Another set of animals received single intraperitoneal treatment of Mcy or oral administration of glibenclamide daily for 30 days as described below. The animals were sacrificed and the tissues were collected and stored at -80°C for molecular and biochemical analyses whereas erythrocytes were isolated and used immediately for insulin binding studies.

Group I: Normal rats treated with buffer [NC].

Group II: Normal rats treated with 1.9mg of Mcy protein/kg bw [NT]

Group III: Diabetic rats treated with buffer [DC].

Group IV: Diabetic rats treated with 1.9mg of Mcy protein/kg bw [DT]

Group IV: Diabetic rats treated with 20mg/kg bw of glibenclamide [DGT]

### Isolation of RNA and Polymerase Chain Reactions

Total RNA was isolated from rat liver tissue using TRI-reagent (Molecular Research Center, Inc., Cincinnati, OH, USA). Briefly, 100 mg of liver was homogenized in 1 ml TRI-reagent and used for cDNA synthesis. Polymerase chain reactions were performed following standard methods using transcript specific primers. Sequences of specific gene primers and the amplicon size were presented in table 1.

**Table 1.**
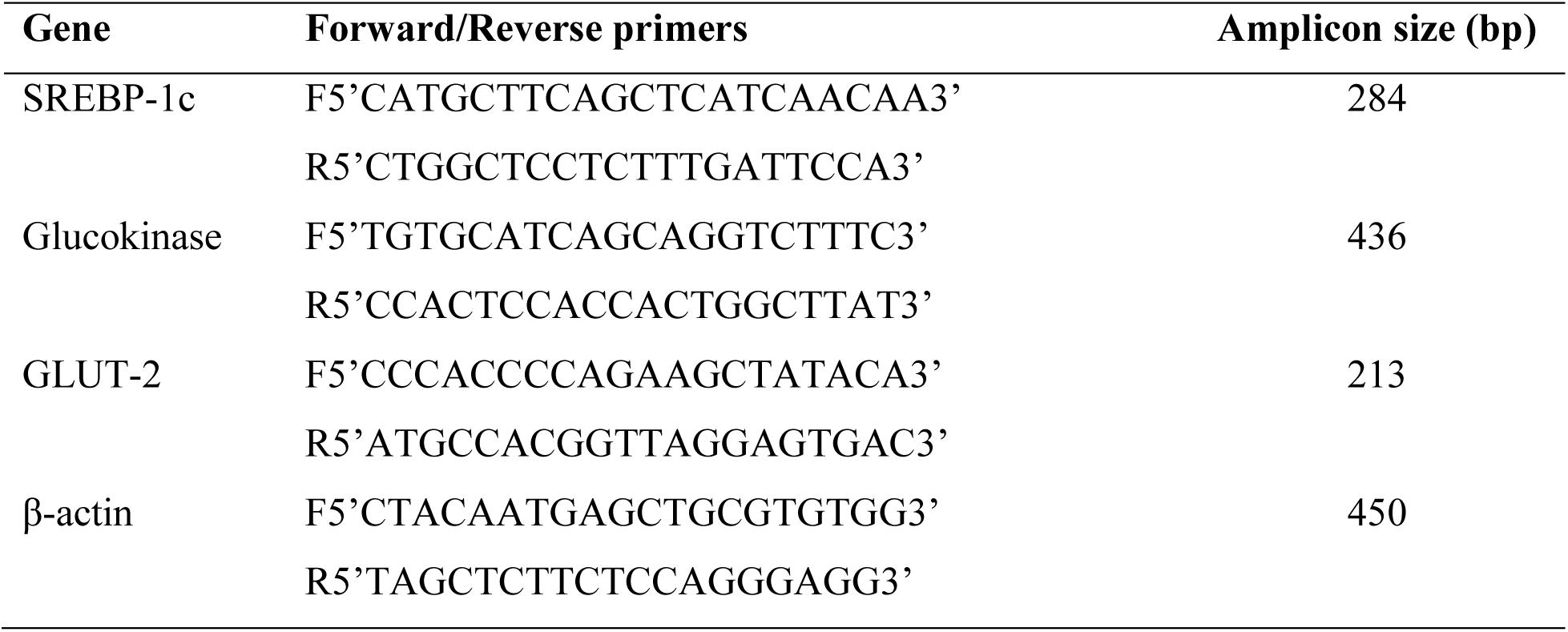
Sequences of the oligonucleotides used in polymerase chain reactions (PCRs)

### Preparation of purified erythrocytes

The receptor assay for erythrocytes was performed following modified methods of Gambhir et al. (1978) Erythrocytes were separated using a percoll density gradient. The erythrocytes were washed thrice with 10 ml of buffer G containing (mmol/L) tris (hydroxymethyl) methylamine, 50; 4-2(2-hydroxyethyl)-1-piperazine ethanesulphonic acid, 50; MgCl_2_.6H_2_O, 10; CaCl_2_, 10; ethylenediaminetetra acetic acid, 2; dextrose, 10; NaCl, 50; KCl, 5; and 1% human serum albumin (pH 7.8) for 10 minutes by centrifugation (4°C, 4500 rpm). On each occasion, the supernatant was removed and the cells were suspended in buffer G and re-spun. After the final wash, the supernatant was removed and the cells were suspended in fresh buffer G containing 1% human serum albumin and used at a cell concentration of 3.94 x 10^9^ cells/ml.

### Binding of ^125-^I to erythrocytes

Erythrocytes (3.94 x 10^9^ cells/ml) were incubated at 15°C with ^125-^I (40 pg in 25 µl) with or without varying amounts of unlabelled insulin (0-0.5 x 10^5^ng) in a total volume of 0.5 ml. After 2.5 h of incubation, duplicate samples were placed in pre-chilled microfuge tubes along with the buffer and dibutylphthalate. Cell-bound and free insulin were separated by centrifugation at 7000 g at 4°C for 10 min. The radioactivity in the cell pellet and supernatant were analyzed in a gamma counter (ECIL, Hyderabad). The data was analyzed by Scatchard analysis (Scatchard, 1949). Specific insulin binding was calculated as the percentage of radioactive insulin bound to 3.94 x 10^9^ cells/ml for erythrocytes. Non-specific binding is defined as the amount of radioactive insulin that remains bound in the presence of 10^5^ ng/ml of unlabeledinsulin. All binding data were corrected for non-specific binding to represent specific cell binding for the purposes of comparison.

The radioactivity bound to the cells was determined by the following

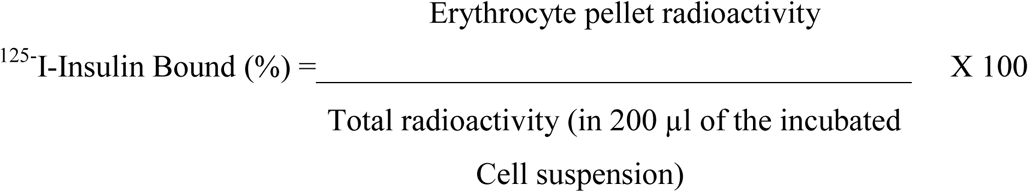

The average number of insulin binding sites per rat erythrocytes was obtained by Scatchard analysis (Scatchard, 1949) of ^125-^I-insulin binding data. Using the amount of the insulin bound (B) and total insulin concentration minus bound (B) as the free insulin concentration (F), a plot of B/F to B was derived.

The average number of estimated sites per cell was calculated using the following expression

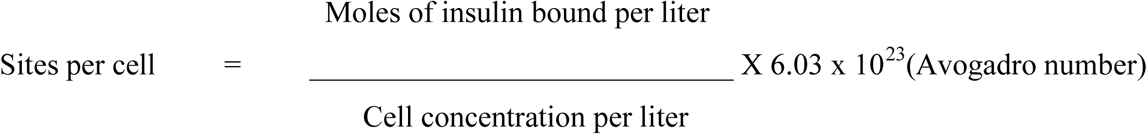

The percentage of specific insulin bound at each concentration of unlabelled insulin was determined by subtracting the percentage of ^125-^I-insulin bound at 1×10^5^ ng per milliliter of unlabelled insulin from the total percentage of ^125-^I-insulin bound at each concentration of unlabelled insulin.

Competitive binding curves were obtained for each erythrocyte suspension and from these, Scatchard plots were drawn to determine the number of insulin receptor sites and receptor affinity.

## Results

The concentration of the protein in ammonium sulfate precipitated fraction was found to be 28mg/ml and the yield was 9% (w/w). The 20% ammonium sulphate precipitated fraction was loaded on the electrophoretic gel along with protein marker. The blot revealed that the Mcy protein (molecular weight 15-17KDa) is detected by insulin antibodies (Fig.1) indicating that Mcy protein potentially shares structural similarity with insulin. We observed two bands recognized by Mcy protein and the higher molecular weight one could be post translationally modified Mcy protein.

### Antihyperglycemic activity of Mcy protein

*In vivo* studies revealed that in normal rats, treatment with Mcy protein (NMT) did not show any hypoglycemic activity (Table 2) whereas treatment of normal rats with insulin (NIT) caused hypoglycemia within 30 min of treatmentas anticipated. In diabetic rats treated with Mcy protein (DMT), the blood glucose levels were significantly reduced up to 79 mg/dl but never showed hypoglycemia (Table 3). However, diabetic rats treated with insulin (DIT) have shown significant hypoglycemic activity.

**Table 2.**
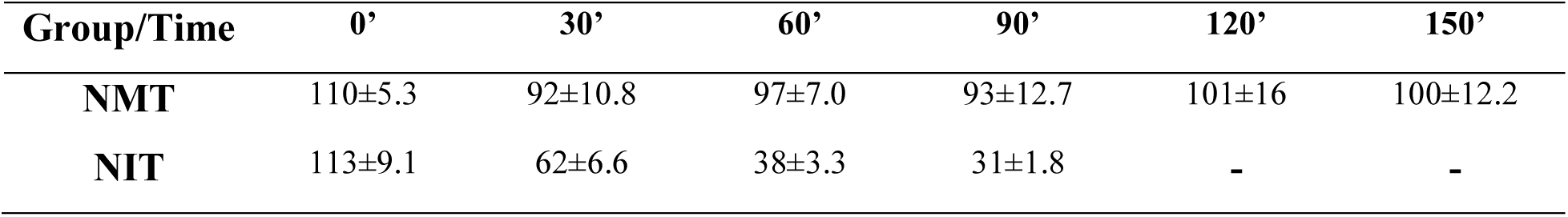
Comparison of glucose lowering activity of Mcy protein versus insulin in normal rats

**Table 3.** Comparison of glucose lowering activity of Mcy protein versus insulin in diabetic rats

| Group/Time | 0' | 30' | 60' | 90' | 120' | 150' | 180' | 210' | 240' |
| --- | --- | --- | --- | --- | --- | --- | --- | --- | --- |
| DMT | 383±7.6 | 341±6.1 | 323±6.4 | 269±4.5 | 218±2.8 | 178±3.8 | 119±6.4 | 101±1.7 | 79±6.3 |
| DIT | 448±1.9 | 296±7.3 | 219±7.8 | 126±7.6 | 84±3.6 | 68±2.1 | 61±1.3 | 50±8.2 | 43±6.1 |

### Gene expression studies

The effect of Mcy protein on hepatic insulin dependent gene transcripts such as GK, GLUT-2 and SREBP-1c from 5 different rat groups were shown in agarose gel electrophoresis (Fig. 2A) and corresponding normalized band intensities were shown in Fig. 2B, 2C and 2D for GK, GLUT2 and SREBP-1c respectively. Semi-quantitative agarose gel (Fig. 2A) also shows the expression of housekeeping gene, β-actin in different groups.

**Figure 1.**
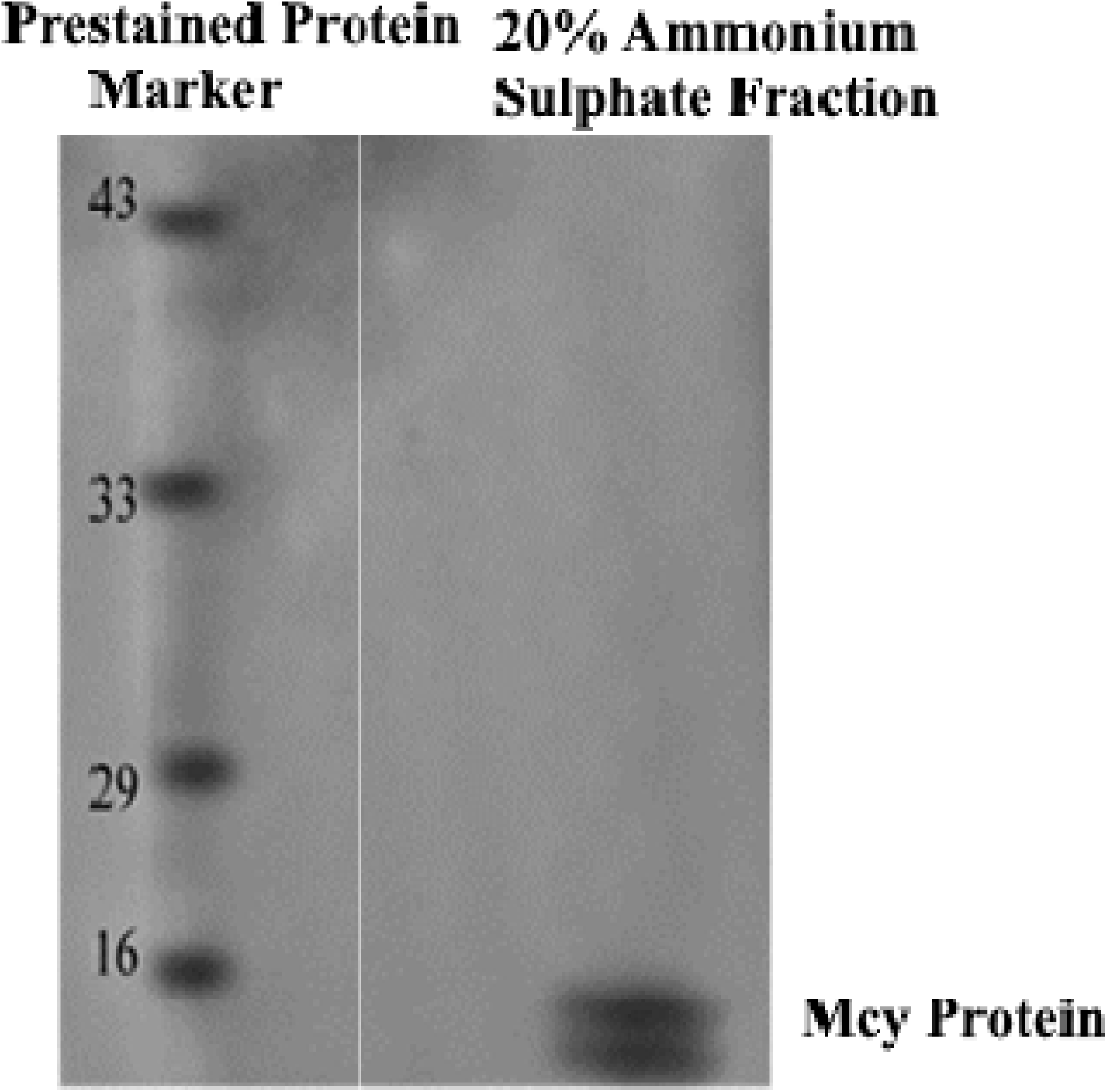
Immunoblot showing recognition of Mcy protein bands from 20% ammonium sulphate precipitated fraction probed with mouse anti-insulin antibody

**Figure 2.**
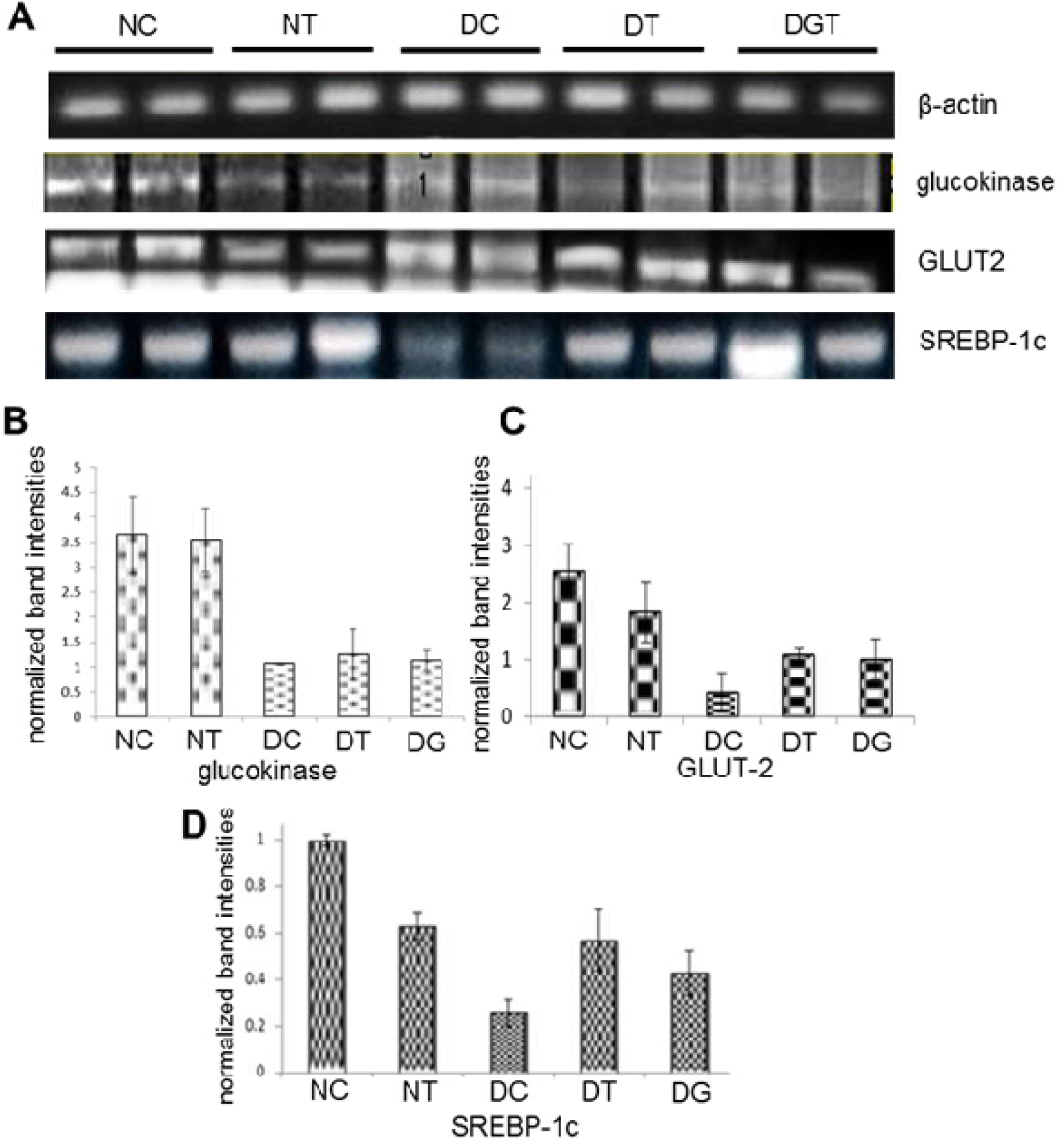
Effect of Mcy protein on the expression the glucokinase (B), GLUT-2 (C) and SREBP-1c (D) gene transcripts normalized to housekeeping gene transcript, β-actin (A).

The expressions of SREBP-1c, GK and GLUT-2 were lower in diabetic group compared to normal group of rats. In Mcy treated diabetic rats there was a significant improvement in the gene expression of SREBP-1c, GK and GLUT-2. The expression of SREBP-1c and GK genes were significantly elevated in diabetic rats treated with Mcy and comparably beneficial than those treated with glibenclamide.

### Receptor binding study

#### Competitive binding curves on the binding of insulin to its receptor on erythrocytes

The ability of non-radioactive insulin to competitively inhibit the binding of ^125^I-insulin to the insulin receptor on cell membranes of erythrocytes in normal, diabetic control and Mcy treated diabetic rats was summarized in **Fig. 3**. Comparison of the plots showed that the insulin receptor on the cell membranes of erythrocytes from diabetic rats bound significantly less ^125^I – insulin than the cells from normal and Mcy treated diabetic rats at the same unlabelled insulin concentration.

**Figure 3.**
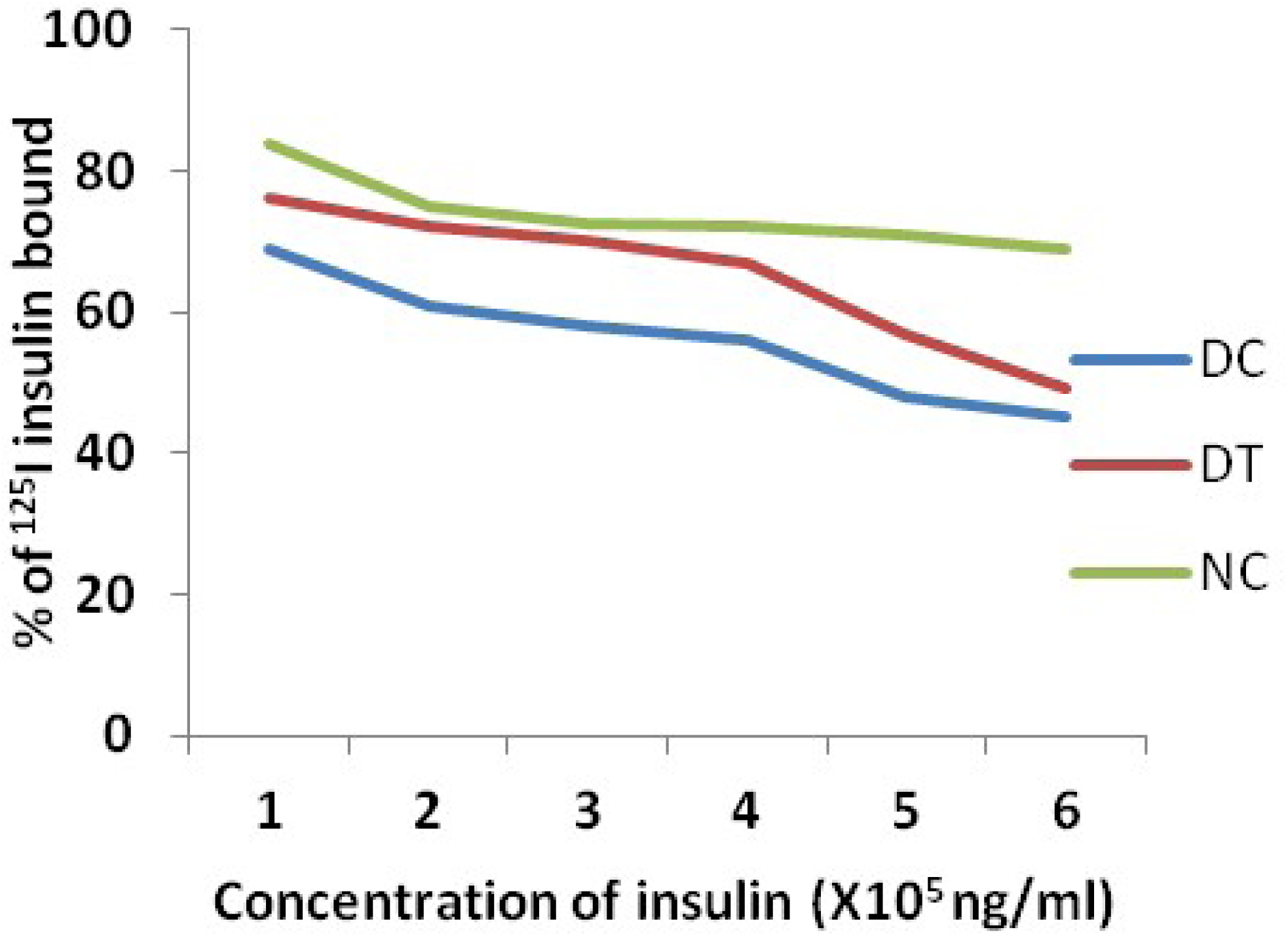
Competitive binding curves showing the effect of Mcy in different groups of rats on the binding of insulin to insulin receptor on erythrocytes. Percentage of ^125^I –insulin bound is plotted as a function of unlabelled insulin concentration. Mean values of 6 rats in each group were considered for the study.

The % ^125^I –insulin bound to the insulin receptor on the cell membrane of erythrocytes from diabetic rats was significantly lower than the percentage of ^125^I –insulin bound to those of normal and Mcy treated diabetic rats at very low unlabelled insulin concentrations (0 to 1ng/ml). Comparison of the competitive curves of the percentage of ^125^I –insulin bound to the insulin receptor on erythrocytes of normal, diabetic and Mcy treated diabetic rats showed slopes that decreased steadily up to an unlabelled insulin concentration of 6 ng/ml.

#### Scatchard plot

The bound/free (B/F) ratio of the labeled hormone is expressed as a function of the bound hormone, yielding a Scatchard plot for erythrocytes. Curvilinear plots were obtained for the diabetic and Mcy treated diabetic rats. A greater B/F ratio implies more bound hormone than free. Comparison of the plots showed that the insulin receptor on the cell membranes of erythrocytes from Mcy treated diabetic and normal rats had higher B/F ratio than that of diabetic rats (Fig. 4). The maximum number of receptors bound by insulin (Bmax) in diabetic control rats and Mcy treated diabetic rats were 150±37.3 and 180±21.1 respectively as calculated from the Scatchard plot.

**Figure 4.**
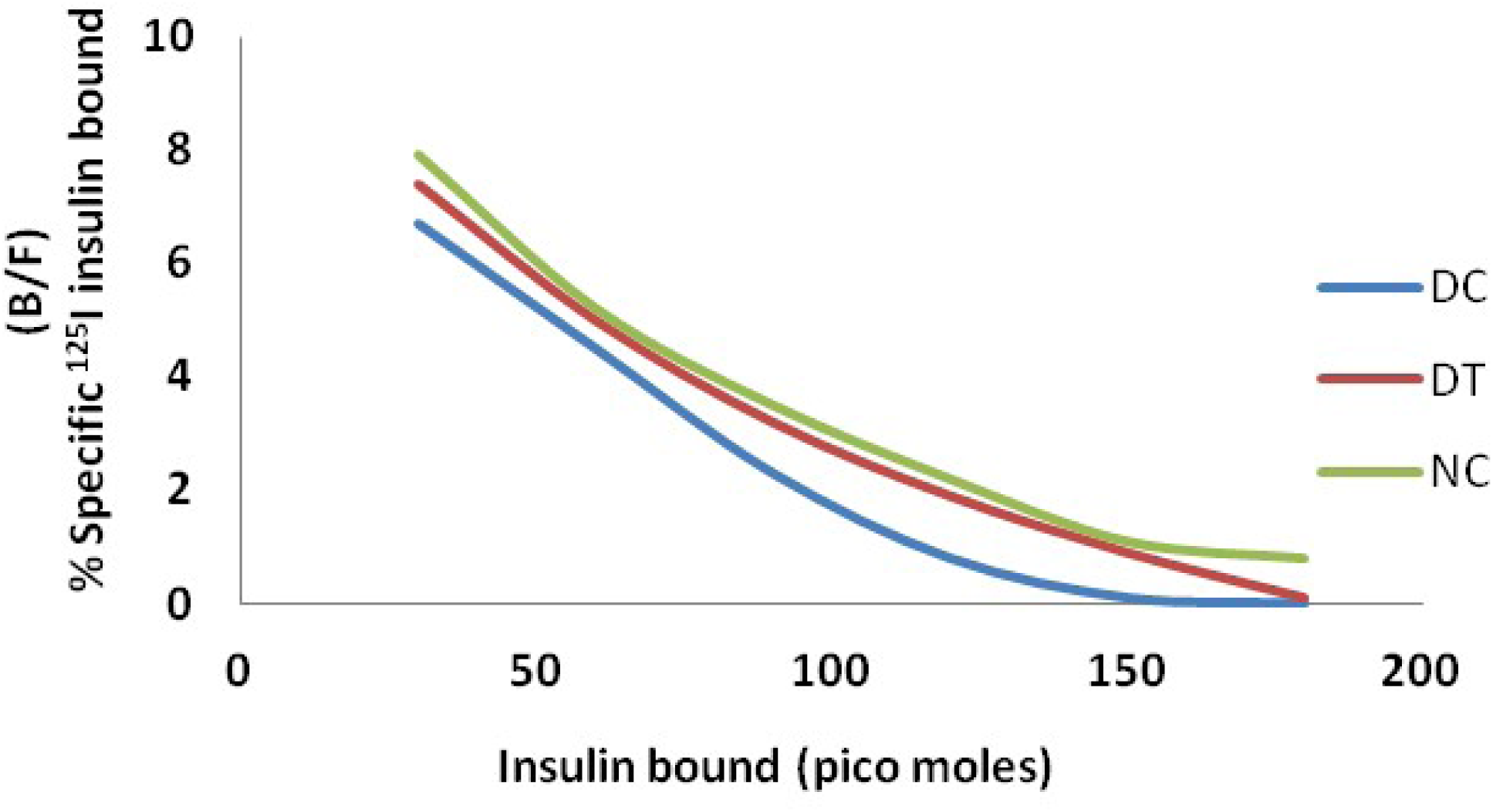
Competition-inhibition studies of Insulin receptor binding Bound and free ratio of ^125^I –insulin (B/F) is plotted as a function of the insulin bound. Extrapolation of the curve to the horizontal axis is used to obtain the total receptor concentration (B_max_) i.e maximum binding.

Scatchard curve for average affinity profile yielded a negative correlation trend between % occupancy 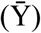 and average affinity (K^−^). In the present study, the highest or “empty site” affinity for Mcy treated erythrocyte was found to be 0.58×10^8^M^-1^ while the average affinity started decreasing when the total sites were occupied i.e 0.57×10^8^M^-1^(Fig. 5).With the increasing occupancy of the receptors by insulin, apparent K‾progressively decreases i.e. there is significant decrease in affinity in the erythrocytes of diabetic rats and increased affinity in rats treated with Mcy protein.

**Figure 5.**
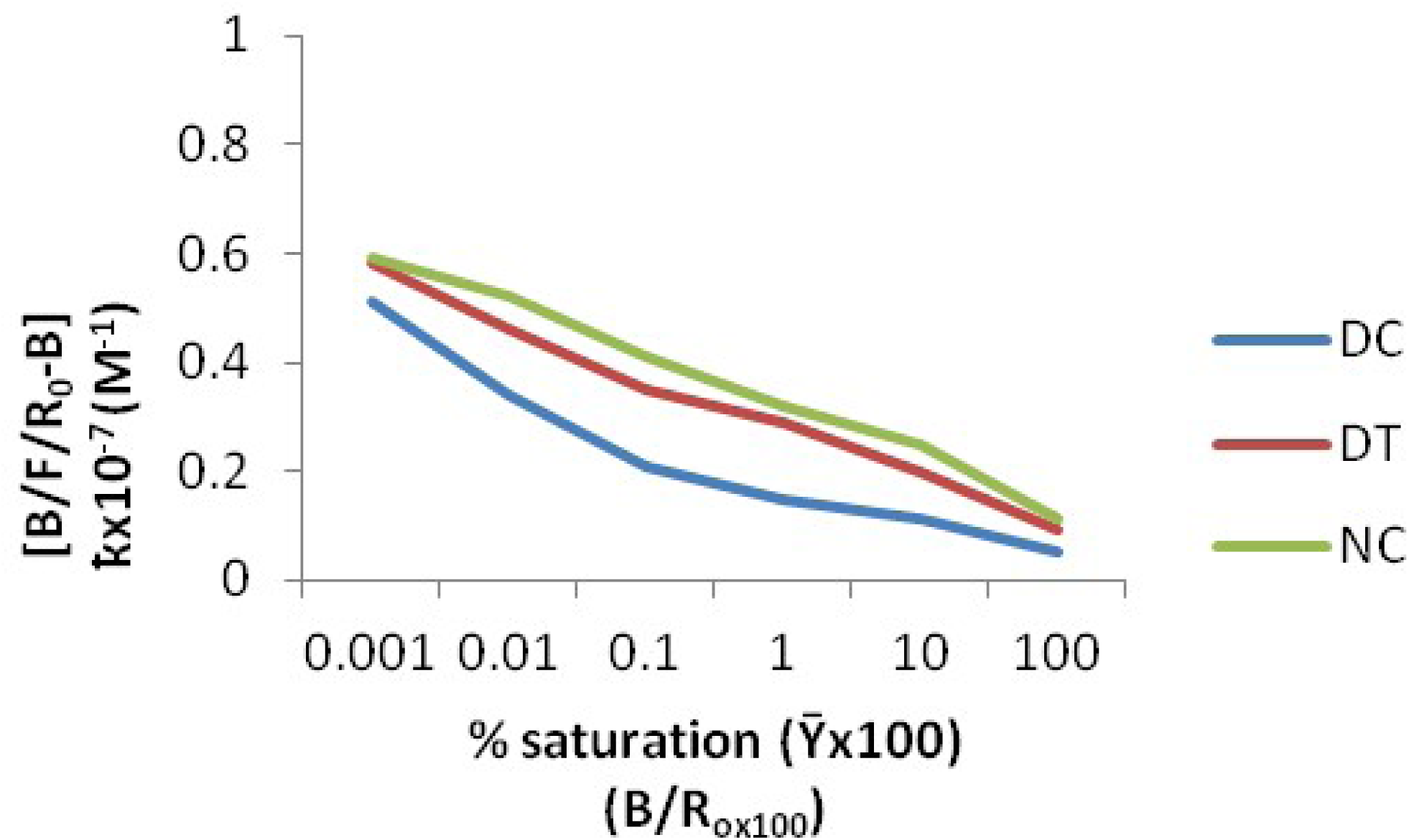
Average affinity binding profile of the ^125^I-insulin

#### Two dimensional gel electrophoresis and MALDI-TOF studies

Isoelectric focusing of the 20% ammonium sulphate precipitated fraction using IEF strip 3-10 pH resulted in separation of different protein spots based on their pI (isoelectric point). Later second dimension gel electrophoresis (SDS-PAGE) resulted in further resolution of the protein mixture according to their molecular weights. The Mcy protein (illustrated in rectangular box) in Fig. 6 aligned between 14 and 18 KDa molecular weight protein markers at pI 5-6.5 as identified earlier was isolated (Rajasekhar et al., 2010) and used for the current studies. This band was excised from the 2D gel after trypsin digestion subjected to MALDI-TOF analysis.

**Figure 6.**
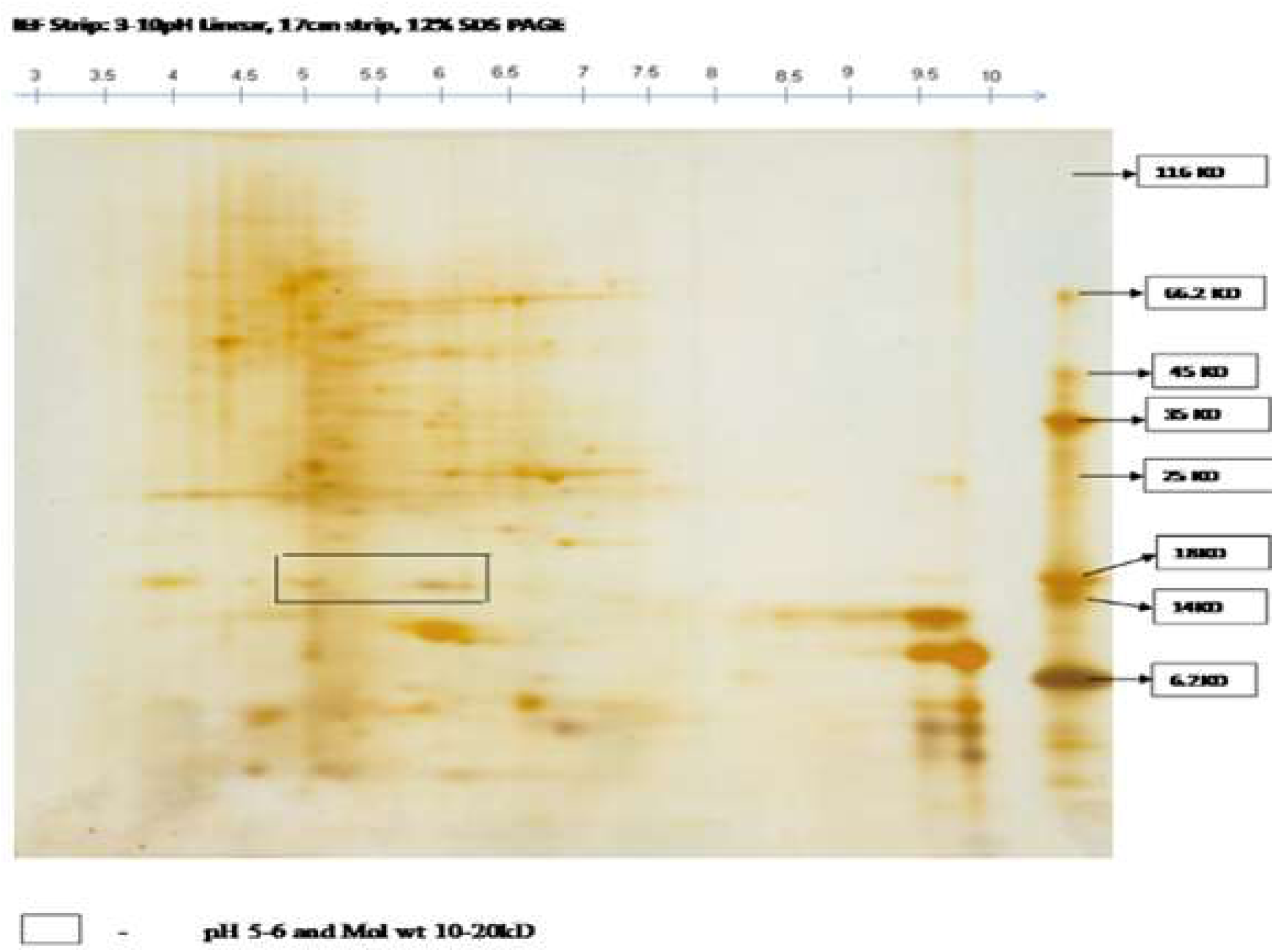
Two dimensional gel electrophoresis of ammonium sulphate fraction of crude protein isolated from the fruits of *Momordica cymbalaria*

Spectra were acquired in the 700–3000 m/z range, processed with Mascot Distiller 2.0.0 (www.matrixscience.com), and the resulting peak lists were used to identify the corresponding proteins in NCBI (non-redundant) and Swiss-Prot databases by peptide mass fingerprinting (PMF) using the Mascot (www.matrixscience.com) search engines. Searches were performed using the following parameters: trypsin as the proteolytic enzyme, allowing for one missed cleavage; carbamido methylation of cysteine as a fixed modification; oxidation of methionine as a variable modification. The mass spectrum obtained for the protein spot is represented in Fig. 7.

**Figure 7.**
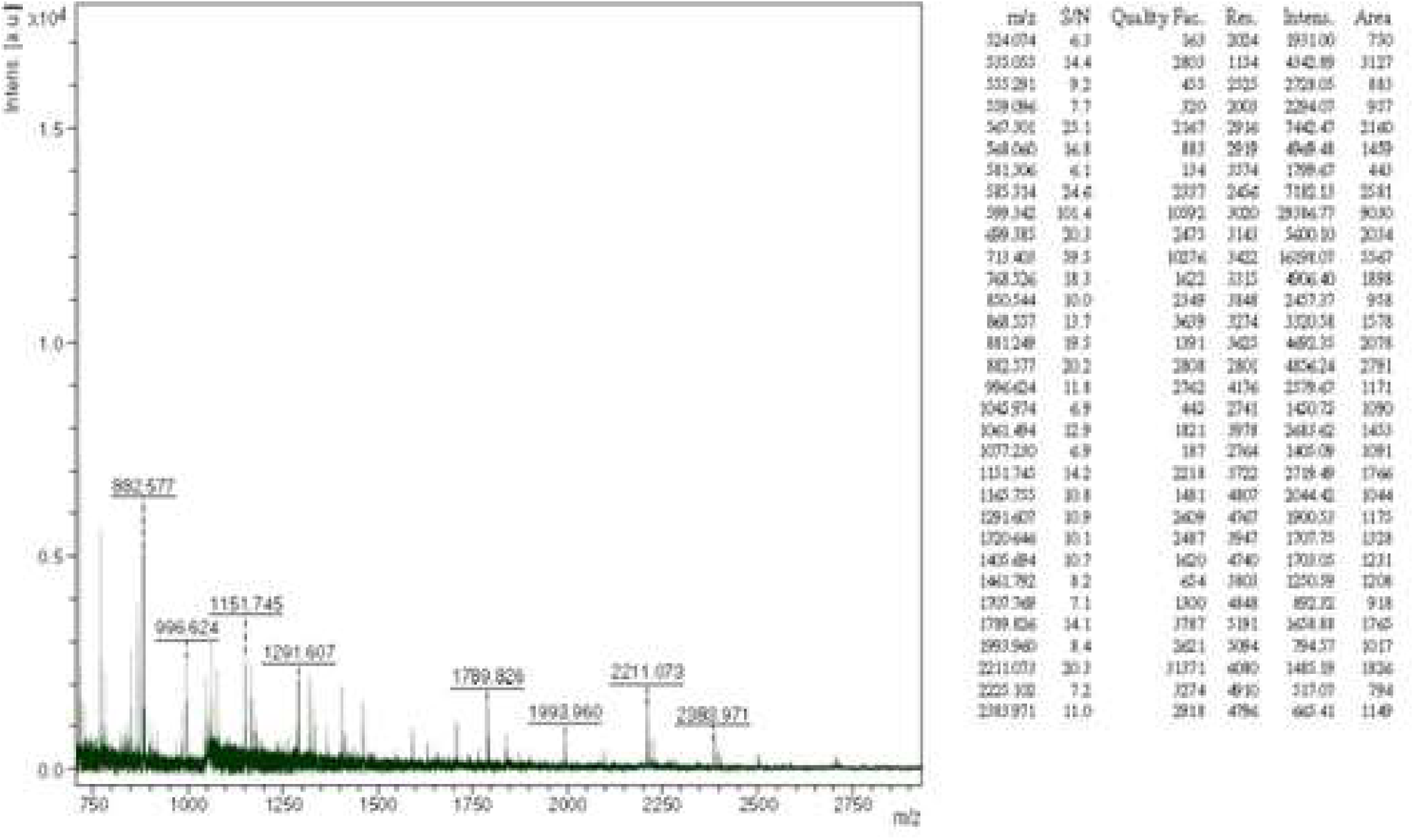
Peptide Mass Finger printing of Mcy protein

## Discussion

Due to presence of high abundant proteins in plants like ribulose-1,5-bisphosphate carboxylase/oxygenase (RuBisCo) or cruciferins, low abundant proteins remain difficult to analyze (Boschetti et al., 2009) Therefore, various fractionation techniques are necessary to enrich these proteins prior to analysis (Kim et al., 2001; Xi et al., 2006). We used 2D gels to separate protein of interest based isoelectric point and molecular weight followed by excision of protein of interest and trypsin digestion for further identification and full characterization of protein/s using mass spectrometry (MS). Peptide mass fingerprinting (PMF) (also known as protein fingerprinting) is an analytical technique for protein identification in which the unknown protein of interest is first cleaved into smaller peptides and identified with a mass spectrometer such as MALDI-TOF or ESI-TOF (Clauser et al., 1991). MALDI-TOF/TOF studies were used to identify the peptide sequence of glycoprotein from *Macrotyloma uniflorum* (Gupta et al., 2013). MS/MS has been effectively used for amino acid sequencing and to investigate the structural information of bioactive peptides derived from different proteins, such as soy protein (Kodera and Nio, 2006) casein (Hernández-Ledesma et al., 2004) and ovotransferin (Majumder and Wu, 2010). In the present study, peptide fingerprint of Mcy matched with that of heat shock protein (HSP)-11, a class I heat shock protein of *Solanum peruvianum*. Small HSPs, HSP25 and HSP27, belongs to α-crystallin family and reported to have neuroprotective and cardioprotective effects against secondary complications of diabetes such as neuropathy, retinopathy, and atherogenesis (Losiewicz and Fort, 2011). HSP22, either expressed at lower levels or its molecular properties are disrupted in the diabetic retina indicating its critical role in diabetic retinopathy (Losiewicz and Fort, 2011; Reddy et al., 2013).

Mcy protein was recognized by anti-insulin antibodies indicating potential structural similarity between Mcy protein and insulin as indicated by immunoblot studies. It is very interesting to note that Mcy protein did not show any hypoglycemic activity in both normal and diabetic rats. Previously insulin like (p-insulin) and non-insulin like (leg insulin) proteins were isolated from plants *Momordica charantia* and soybean respectively (Khanna et al., 1976; Watanabe et al., 1994). Further, based on sequence similarities Watanabe et al. reported that this leg insulin is also present in pea seeds, adzuki beans, mung beans, and carrot (Watanbe et al., 1994). Collip (1923) reported the presence of insulin-like material known as glucokinin that is diverse in function compared to insulin.

Sterol regulatory element binding proteins (SREBPs) belongs to helix-loop-helix leucine zipper family of transcription factors that regulates fatty acid & cholesterol synthesis (Brown and Goldstein, 1997, 1999). Three isoforms of SREBPs are known to date SREBP-1a, SREBP-1c & SREBP-2. SREBP-1c plays a crucial role in regulating gene expression of lipogenic enzymes (Shimano, 2000, 2001; Horton et al., 2002). Insulin is known to be a trigger for enhancing the SREBP-1c transcription levels in livers of STZ induced diabetic rats (Shimomura et al., 1999). In the present study SREBP-1c expression is down regulated in diabetic rats which may be due to insulin deficiency. Whereas elevated levels of SREBP-1c expression in Mcy treated diabetic animals to near normal could be due to elevated insulin levels (Marella et al., 2015) or insulin mimicking activity of Mcy protein. Direct action of Mcy protein on SREBP-1c gene transcription cannot be ignored.

GLUT-2 is membrane protein, transports glucose across the hepatic plasma membrane bi directionally. GLUT-2 is mainly expressed in the liver, β-cells of the pancreas and basolateral membrane of kidney proximal tubules and plays an important role in glucose homeostasis. GLUT-2 expression is also regulated by SREBP-1c levels. Additionally, insulin resistance also plays a central role in glucose transport. Increased levels of SREBP-1c trigger the up regulation of GLUT-2 and vice versa (Im et al., 2005). In our study GLUT-2 gene expression is down regulated in STZ induced diabetic rats and it could be due to low insulin levels that possibly reduced the expression of SREBP-1c which in turn diminished GLUT-2 expression. Increased insulin levels may also be contributing for maintaining normal GLUT-2 transcript levels in Mcy treated diabetic animals through alternate pathways.

Hepatic glucokinase is a crucial enzyme in maintaining glucose homeostasis. Glucokinase activity in liver is low during fasting and in diabetes mellitus. Physiologically, a low glucokinase activity favors the release of glucose into the blood circulation via gluconeogenesis. Conversely, a high glucokinase activity promotes glycogen deposition in the liver (Hers and Hue, 1983). The level of hepatic glucokinase activity appears to be determined essentially by regulation of the rate of enzyme synthesis, with insulin playing a leading role as an inducer (Weinhouse, 1976). Foretz et al. (1999) stated that the transcriptional activity of SREBP-1c is required for insulin action on the glucokinase gene expression. The over expression of the dominant positive form of SREBP-1c mimics the effects of insulin on glucokinase gene. Further, in STZ diabetic mice, adenovirus-mediated over-expression of SREBP-1c resulted in the increase of hepatic glucokinase expression and activity along with an increase in the hepatic glycogen content (Foretz et al., 1999) mimicking the effect of insulin. The expression of GK gene was significantly increased in Mcy treated diabetic rats which could be due to increased SREBP-1c gene expression. Our studies indicate that the normalized levels of glucokinase, GLUT-2 could be due to elevated levels of SREBP-1c that is essential for glucokinase expression and insulin action. The beneficial effect of Mcy protein treatment on the expression of the key regulatory genes investigated in our present studies are comparable to the treatment of glibenclamide.

Activation of insulin receptors lead to cellular mechanisms that directly affect glucose uptake by regulating the number and function of protein molecules in the cell membrane that facilitate glucose transport into the cell. Insulin binds to the extracellular portion of the alpha subunits of the insulin receptor. This, in turn, causes a conformational change in the insulin receptor that activates the intracellular kinase domain of the beta subunits. Auto phosphorylation and dephosphorylation in turn regulate the activity of kinase domain of the receptor as well as tyrosine residues in the IRS-1 protein. Further, insulin action is alsoregulated by the number of receptors on the plasma membrane, a decrease in the amount of receptors also leads to reduced or termination of insulin signaling.

With the introduction of the human erythrocyte insulin receptor assay by Gambhir et al. (1977) erythrocytes were shown to possess insulin binding characteristics similar to monocytes and other human and animal tissues. Using erythrocytes, Robinson et al. (1979) showed decreased insulin binding in diabetics, and Gambhir et al. (1977) indicated a defect in insulin binding in uremic patients. Many investigations suggested the effect of plant and plant products on insulin receptor binding to erythrocytes (Murugan et al., 2008). Pari et al. (2004) demonstrated the influence of *Cassia auriculata* flowers on insulin receptors in streptozotocin induced diabetic rats. In the present study, treatment with Mcy showed significant antihyperglycemic activity and this may be due to insulin secretagogue potential of Mcy protein to release insulin from the existing β-cells of pancreas. The hyperglycemia in untreated diabetic rats could be partially due to a significant decrease in the receptor concentration per cell and also due to a marginal decrease in the affinity of receptors to insulin. Mcy protein has been shown to potentiate insulin secretion causing a significant decrease in blood glucose which in this study further supported by improved number of insulin binding sites with the administration of Mcy protein (Rajasekhar et al., 2010). In the present study, we also showed that there is a decrease in the binding affinity of insulin to insulin receptors in diabetic rats which was significantly enhanced with Mcy treatment indicating the beneficial role of Mcy protein in increasing insulin receptors as well as affinity of insulin to insulin receptors.

## Conclusion

The MALDI-TOF analysis through MASCOT search revealed similarity of the Mcy protein with heat shock protein (HSP 19.9) of *Solanum peruvianum*. The small HSPs were known to have neuroprotective and cardioprotective effects which are in support of antidiabetic functions of Mcy protein and its beneficial role in preventing secondary complications of diabetes. Mcy protein did not show hypoglycemia in both normal and diabetic animals. Further, our studies demonstrate that Mcy protein treatment enhanced the number of insulin receptors and affinity of insulin with its receptor binding in addition to its significant beneficial role on the levels of glucokinase, GLUT-2 and SREBP-1c, key carbohydrate and lipid metabolism regulators.

## Acknowledgements

This work was supported by University Grants Commission (to CA, major research project 37-360/2009), New Delhi, India. The first author thanks Dr. Adinarayana, University of Hyderabad and Dr. E.G.T.V. Kumar for their help in Insulin studies.

